# Prefrontal Cortex Encodes Value Pop-out in Visual Search

**DOI:** 10.1101/2023.01.27.525832

**Authors:** Mojtaba Abbaszadeh, Armin Panjehpour, Mohammad Amin Alemohammad, Ali Ghavampour, Ali Ghazizadeh

## Abstract

Recent evidence shows that long-term object value association can lead to efficient visual search. However, the neural mechanism of this value pop-out has yet to be understood. Given the known role of the ventrolateral prefrontal cortex (vlPFC) in visual search and value memory, we recorded its single-unit activity (n=526) in two macaque monkeys while they engaged in the value-driven search. Monkeys had to determine whether a high-value target was present within a variable number of low-value objects. Interestingly, differential neural firing, as well as gamma-band power, indicated the presence of a target within ∼150ms of display onset. This differential activity was negatively correlated with search time and became less display size-dependent for more efficient searches. On the other hand, neural firing and its variability were higher in inefficient searches. These findings reveal the neural code within vlPFC for rapid detection of valuable targets, which can be crucial for animals faced with competition.

**Significance Statement:** Search for rewarding objects is ubiquitous and crucial for animals and humans alike. Up until recently, it was thought that visual search for valuable targets that were otherwise not distinct by low-level features should be serial and slow. Contrary to this belief, we showed that given sufficient reward training, valuable objects can be found efficiently in search suggesting a value pop-out neural mechanism. Importantly, we reveal the neural activity in the prefrontal cortex (PFC) to be predictive of the degree of value-driven search efficiency. Given the role of PFC in object value memory, these results show how PFC can translate such memories to emulate the parallel processing of visual information independent of low-level visual features.

## Introduction

Rapid detection and acquisition of valuable objects is an important skill for animal survival. Nevertheless, when valuable objects are surrounded by and are not easily distinguishable from other low-value or irrelevant objects, the search becomes inefficient and slow. For a long time, it was believed that the guiding features that can support efficient search are only limited to low-level visual features such as color and orientation (Wolfe, 1994; Wolfe & Horowitz, 2004). A recent study showed that a high-level feature such as object value draws automatic gaze (Ghazizadeh, Griggs, & Hikosaka, 2016a) and can support parallel search provided that object reward association was overtrained (Ghazizadeh, Griggs, & Hikosaka, 2016b). Given the fact that the neural circuitry in the early visual cortices is not known to process value the same way they do orientation, color or size (Gur, Kagan, & Snodderly, 2005; Kobatake & Tanaka, 1994; Maunsell & Treue, 2006), the neural circuitry that can support the value pop-out in visual search remained unknown.

Single unit recordings as well as fMRI, revealed a key role for vlPFC in learning and maintaining object value memories, especially for those overtrained with reward (Ghazizadeh, Griggs, Leopold, & Hikosaka, 2018; Ghazizadeh & Hikosaka, 2021; Ghazizadeh, Hong, & Hikosaka, 2018). Furthermore, vlPFC is shown to be part of the circuitry that guides gaze based on object reward history, which includes superior colliculus (SC, Griggs et al., 2017) and basal ganglia nuclei such as caudate (Kim, Amita, & Hikosaka, 2017) and substantia nigra reticula (SNr, Hikosaka, Kim, Yasuda, & Yamamoto, 2014). In addition, previous studies have implicated vlPFC in visual search (Bichot, Heard, DeGennaro, & Desimone, 2015; Bichot, Xu, Ghadooshahy, Williams, & Desimone, 2019). Inactivation of vlPFC neural regions decreased subjects’ efficiency in finding the target in a feature-based visual search task.

Based on these findings, an important question arises as to whether and how fast the neurons in vlPFC can discriminate between trials that do and do not contain the high-value objects (target-present and target-absent trials, respectively) and whether those responses are implicated in the efficiency of value-oriented search. To address these questions, the response of vlPFC neurons was recorded while monkeys engaged in efficient and inefficient value-driven search with fractal objects that had a variable duration of past reward training. Notably, we find a rapid and robust differentiation of target-present and target-absent conditions in both neural firing and local field potential in vlPFC. This differentiation was much more prominent and less display size-dependent in more efficient searches.

## Results

To understand the role of vlPFC in efficient visual search, the response of well-isolated neurons was recorded while macaque monkeys performed a value-driven search task (see methods). The values of fractals were learned prior to the search task in an object-value training task that involved saccading to single objects to receive their associated reward (value training, Fig. 1A-B). Each value training was done with 8 fractals, half of which were associated with low reward and the other half with high reward (bad and good fractals, respectively). Objects sets were trained either for 1-day or >5 days (over-trained fractals). Performance during choice trials that were interspersed during the value training showed that for 1-day trained fractals subjects chose good objects nearly perfectly and well above chance (∼95%), similar to the choice performance in over-trained fractals, which albeit were slightly but significantly higher (∼98%, Fig. 1C) suggesting nearly perfect knowledge of object values in both fractal groups. The target acquisition time defined as the time from the go cue (fixation offset) until landing on the target, was also similar between 1-day trained and over-trained fractals for both good and bad fractals (ps > 0.05). As expected, the acquisition time for the good fractals was significantly lower than the bad fractals in both training groups (t_238_ = 7.93, p < 0.0001), consistent with previous literature (Kim, Ghazizadeh, & Hikosaka, 2015).

**Fig 1.**
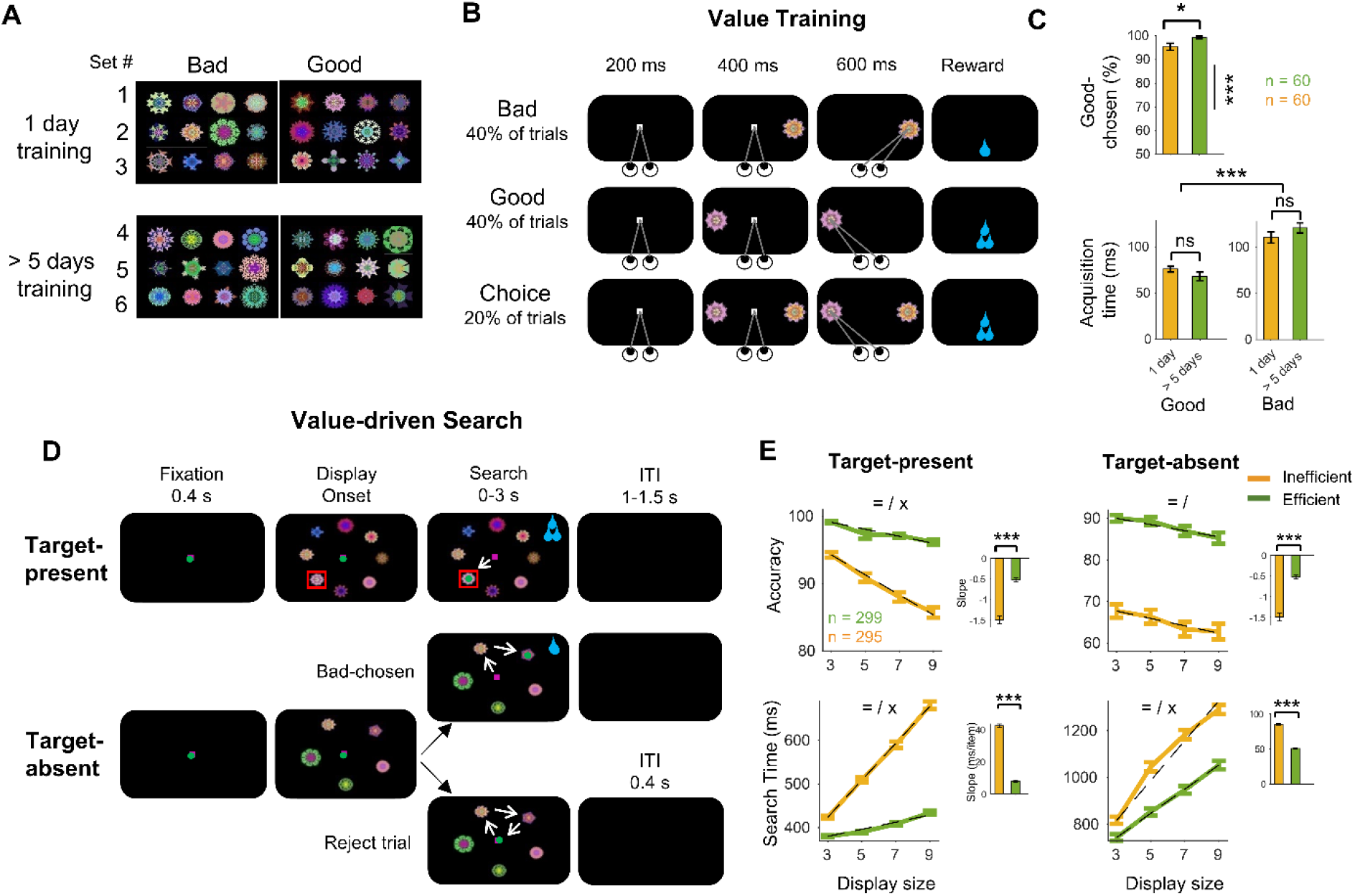
Stimuli and experimental paradigm. A) Fractal objects were trained in sets of eight, four big-reward (good) and four small-reward (bad) fractals. Monkeys S and H were trained with 95 and 92 fractal sets, respectively. Fractals set were value trained for one day or more than five days (> 5 days). B) Value training: this task consisted of forced and choice trials. In the forced trials, one fractal (good or bad) was randomly presented in one of eight locations on the screen after a central fixation, and the monkey made a saccade to the fractal to receive the reward. In the choice trials, two fractals (good and bad) were randomly presented in two of eight locations on the screen after a central fixation, and the monkey chose one to receive the corresponding reward. C) Behavioral performance of subjects during the value training task. Precent of good object chosen in choice trials (top) and saccade acquisition time in force trials for good objects (left bottom) and bad objects (right bottom) for the one day >5 day trained objects during value training task. D) Search task: this task consisted of target present (TP) trials (top) and target absent (TA) trials (bottom). In each trial, 3, 5, 7, or 9 fractals were simultaneously presented equidistant on an imaginary circle. In TP trials, only one fractal is good and in TA trials, all fractals were bad. The monkey could get big reward by finding the good fractal and gazing at it or reject the trial by fixating the center to get to the next trial rapidly. Choosing a bad object resulted in small reward. E) Behavioral performance of subjects during search task. Top panels show percent accuracy of finding good objects across display size in TP trials (left) and of rejecting the trial in TA trials (right) for efficient (green) and inefficient (orange) search sessions across display size (linear fit is shown by dashed line). Inset bar plots show average accuracy slope of linear fit for both searches. Bottom panels same format as top panels but for the search time to find good object or reject the trial. (=, /, x: main effect of search efficiency, display size, and interaction, respectively; *p < 0.05, **p < 0.01, ***p < 0.001)

### Over-training with reward creates efficient search

During the search task, monkeys had to find a single good fractal surrounded by a variable number of bad fractals in target-present (TP) trials to receive high reward and had to reject the trial by coming back to the center in target-absent (TA) trials in which all displayed objects were bad. Rejection of a TA trial resulted in a quick procession to the next trial (shortened ITI, Fig. 1D). The monkey would receive low reward if he chose a bad fractal and would receive high reward if he found good object in TP trials. Previously, we found that despite good knowledge of value in both groups of fractals in choice trials, only the search with over-trained fractals tends to be efficient (Ghazizadeh et al., 2016b). Consistently, here our results showed that the TP search times and search slopes (search per item of display size) for overtrained sets were significantly smaller than the slopes for 1-day trained sets (ts < -8.3, ps < 0.0001), suggesting the emergence of the efficient search for fractals that were overtrained with value (Supp Fig 1A).

Behavioral search slopes in both monkeys showed a wide range from inefficient search with slopes >40 ms/item to efficient search with slopes near 0 ms/item in TP trials in both monkeys (Supp Fig 1B, Monkey H TP slopes were overall smaller than Monkey S, t_532_= 3.22, p = 0.001). We grouped searches into efficient and inefficient types by considering slopes in the lower 40 percentile and higher 60 percentile based on each monkey slope distribution.

While the accuracy of good object choice in TP trials was higher than 85% in inefficient search, this was significantly lower than the 95% accuracy in the efficient search across all display sizes (Efficiency main effect: F_1,2336_ = 498.88, p < 0.0001, Fig. 1E). Notably, the accuracy of TP trials in the inefficient search was more display size-dependent compared to efficient search and rapidly fell across all DS from about 95% in 3 object DS to about 85% in 9 object DS for inefficient searches (interaction: F_3,2336_ = 15.54, p < 0.0001, DS main effect: F_3,2330_ = 45.29, p < 0.0001, Fig. 1E). A similar degradation of accuracy in rejecting TA trials in inefficient vs efficient search was also observed (Efficiency main effect: F_1,2330_ = 568.72, p < 0.0001). Although here, DS dependence for the inefficient search was higher than efficient search (F_3,2330_ = 4.58, p = 0.003, Fig. 1E), the interaction of efficiency and DS was not significant (F_3,2330_ = 0.042, p = 0.98, Fig. 1E).

Search time was significantly higher in efficient vs inefficient search in both TP (F_1,2336_ = 1923, p < 0.0001) and TA trials (F_1,2330_ = 271, p < 0.0001) and it increased significantly with DS (DS main effect: F_3,2330_ > 182, p < 0.0.0001). There was a significant interaction between search type and DS with lower slopes in the efficient search (F_3,2336_ > 170.04, p < 0.0001, Fig. 1E). This was expected in TP trials by the definition of efficient and inefficient search based on TP search slopes but was nontrivial for TA trials.

### High value signal independent of DS in vlPFC is concurrent with efficient search

To examine involvement of vlPFC in value-driven search, the responses of single neurons were recorded and compared during the efficient and inefficient search types. Fig. 2 shows two example vlPFC neurons with responses time-locked to display onset across different display sizes during the efficient and inefficient searches (two-search neurons). In both search types, both neurons showed a clear visual response to display onset. One of the neurons fired more strongly to TP compared to TA trials (TP-pref neuron, Monkey H: Neuron# 559, Fig 2A), while the other fired more strongly to TA compared to TP trials (TA-pref neuron, Monkey S: Neuron# 220, Fig 2B). In both neurons, the differential firing which started as early as ∼100ms was much more pronounced in the efficient search compared to the inefficient search was due to the presence or absence of the good object and will be referred to as the value signal.

**Fig 2.**
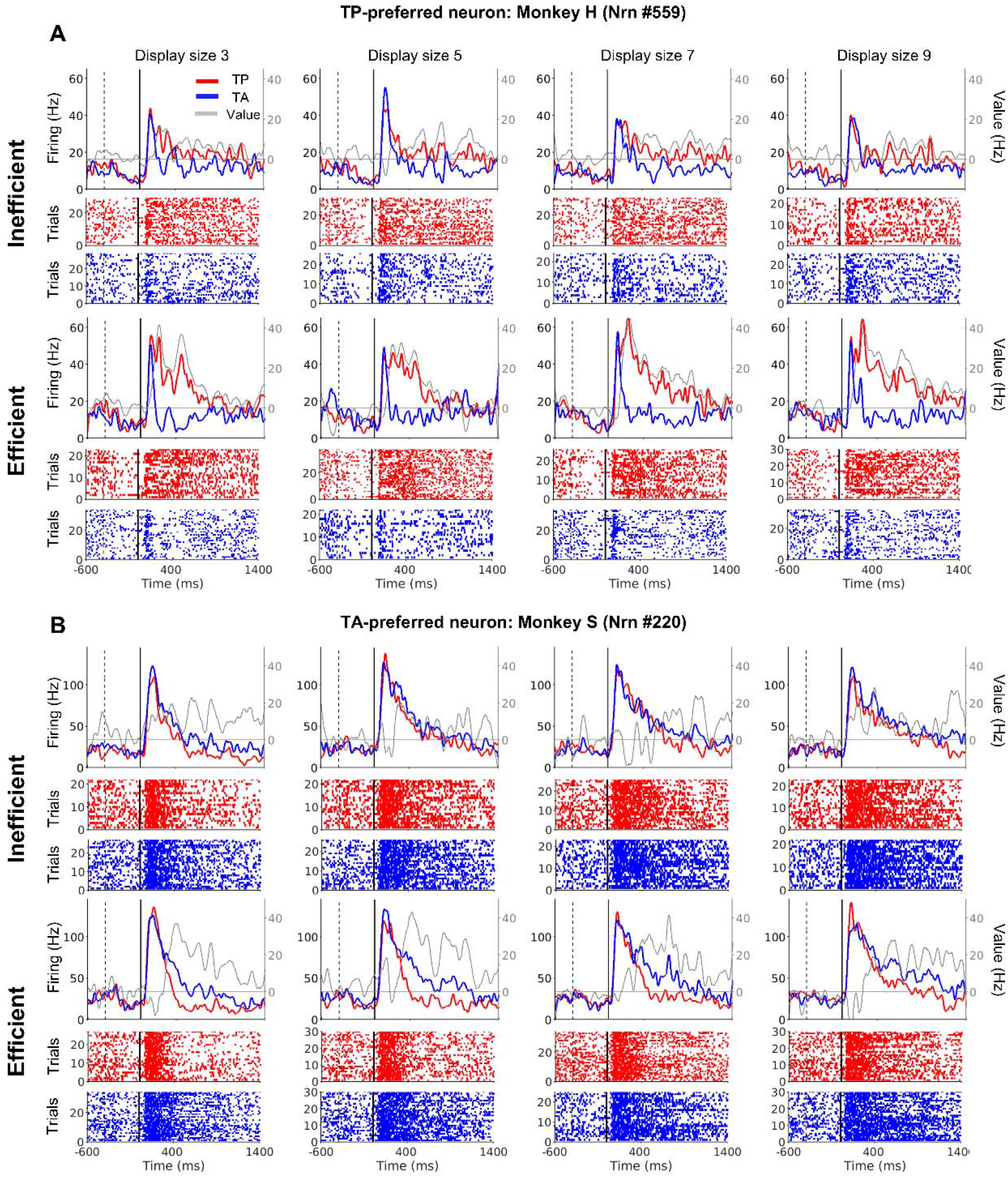
Response of two vlPFC example neurons during efficient and inefficient searches. A) The response of a TP-preferred example neuron during the inefficient and efficient searches. The peristimulus time histograms (PSTH, first row) and raster plots (second and third rows) time-locked to the display onset of for target-present (TP, red) and target-absent (TA, blue) trials across display sizes 3, 5, 7, and 9, respectively, in the inefficient (top) and efficient (bottom) searches. Black line shows TP minus TA firing. B) Same format as A, but for a TA-preferred example neuron. Black line shows TA minus TP firing. Dashed and solid lines on each plot indicate the fixation onset and display onset time, respectively.

To determine the TP and TA discriminability across all recorded neurons, the area under the receiver operating characteristic curve was calculated for each neuron (value AUC). Value AUCs above 0.5 indicate higher firing in TP compared to TA trials (TP-pref), and below 0.5 indicates higher firing in TA compared to TP trials (TA-pref). In both search types, the average of value AUC across the population was significantly above 0.5 (efficient: t_292_ = 5.98, p < 0.0001; inefficient: t_296_ = 4.84, p < 0.0001), suggesting a shift toward TP preference with a larger percentage of vlPFC neurons showing significantly TP-preference (Supp Fig 3A; efficient: X^2^ = 25.99, p < 0.0001; inefficient: X^2^= 5.65, p = 0.01). The percentage of neurons with no significant value AUC (NS neurons) was higher in inefficient search (X^2^ = 11.18, p = 0.0008). Supp Fig 3B shows the average response of all types of neurons (TP-pref, TA-pref, and NS) for efficient (top) and efficient (bottom) collapsed over DS.

**Fig 3.**
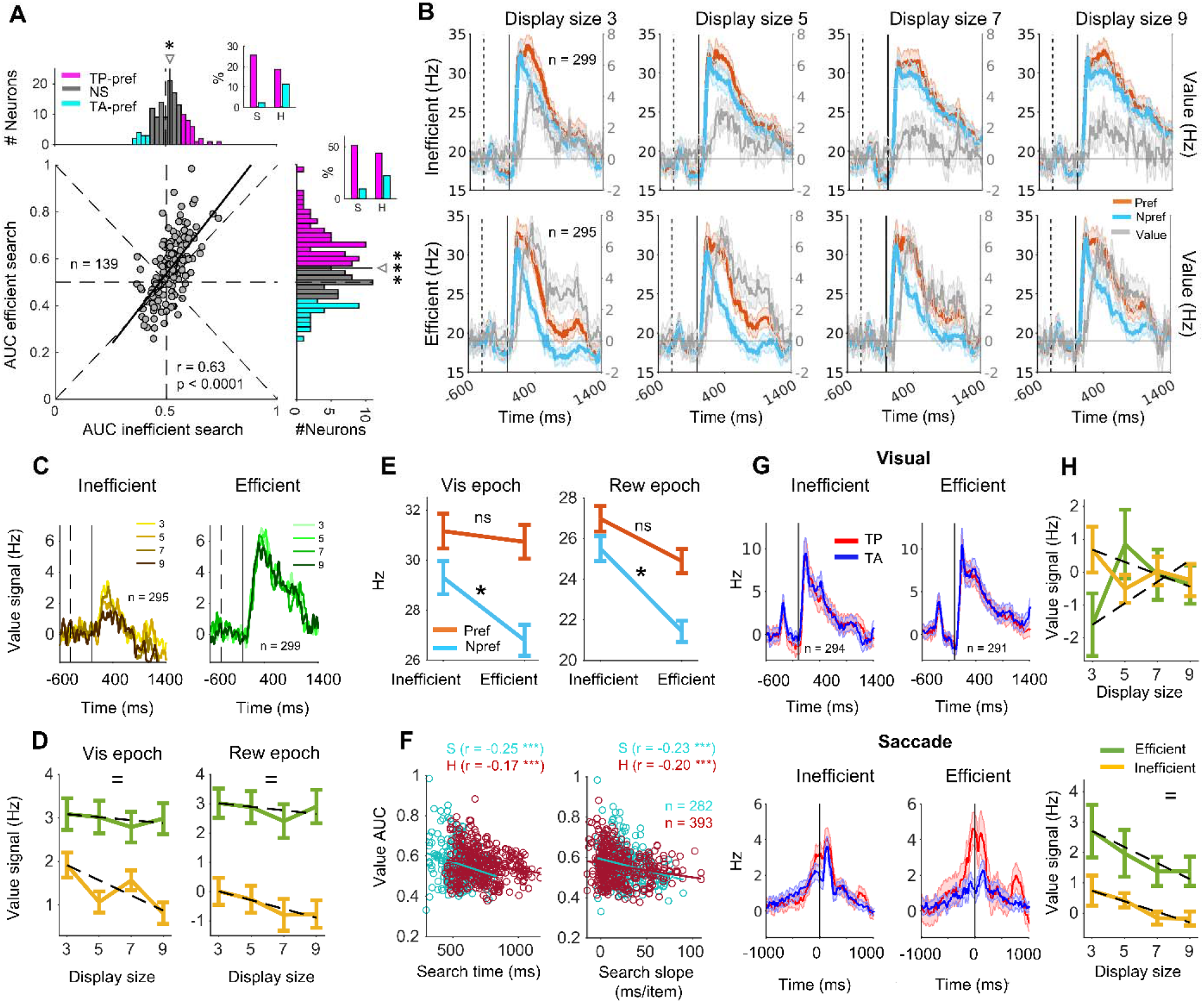
Value signal of vlPFC neurons in inefficient and efficient searches. A) Scatter plot of value AUC in the inefficient and efficient search for two search neurons along with marginal histograms. The solid dark line indicates the linear fit (Deming regression). The pink, cyan, and gray colors in the value AUC histograms show the TP-preferred (TP-pref, significant AUC > 0.5), the TA-preferred (TA-pref, significant AUC < 0.5), and nonsignificant (NS) neurons, respectively. Insets indicate the percentage of TP- and TA-pref neurons for both monkeys in the search types. B) Population average of PSTH for each display size across neurons for the inefficient search (top row) and the efficient search (bottom row). Red and light blue show preferred and non-preferred values, respectively. Black shows the difference. Dash and solid lines same as Fig 2. C) The difference between preferred and non-preferred signals (value signal) for the inefficient (left) and the efficient (right) searches for all display sizes. D) The average value signal across neurons during visual epoch (100-400ms relative to display onset, left) and reward epoch (800-1100ms relative to display onset, right), respectively, across the display size for both the inefficient (orange) and the efficient (green) searches. The dashed lines show the linear fit of the value signal as a function of display size. E) Average neural firing for preferred (red) and non-preferred (light blue) signals across search types for visual (left) and reward (right) epochs. F) Scatters show the correlation of value AUC of neurons in a given session with search time (left) and search time slope (right) in TP trials separately for monkey S (cyan) and monkey H (dark red). G) Deconvolution of population average of visual (top) and saccadic (bottom) parts of neural response for TP (red) and TA (blue) trials in the inefficient (left) and efficient (left) searches. H) Value signal for the visual (100-400ms relative to display onset, top) and saccadic (-150-150ms relative to saccade onset, bottom) part of deconvolved neural responses in the inefficient (orange) and the efficient (green) searches. (=, /, x: main effect of search efficiency, display size, and interaction, respectively; *p < 0.05, **p < 0.01, ***p < 0.001)

Using two-search neurons there was a significant positive correlation between value AUCs between the two search types (r = 0.63, p < 0.0001, Fig 3A). This meant that neurons had consistent TP or TA preference across search types and the ones with strong value signal in one search type tended to have strong value signal in the other search type. Importantly, the slope of linear relation between efficient vs inefficient search was significantly larger than one (100 resamplings, t_99_ = 49.01, p < 0.0001), suggesting an enhancement of value signal within two search neurons in the efficient search.

One may quantify the population activity in search by averaging responses to preferred and non-preferred trial types (TP or TA) using cross-validation (Supp Fig 3C, see methods). The average preferred versus non-preferred responses across all neurons revealed stronger differentiation between preferred and nonpreferred values and a higher value signal in the efficient compared to inefficient search (Fig 3B-C). The onset of value signal could be detected as early as 146±4.46 ms after display onset across neurons. The main effect of the search type on the value signal was significant for visual (F_1,2364_ = 55.9, p < 0.0001) and reward (F_1,2364_ = 78.9, p < 0.0001) epochs (Fig 3D). Notably, while value signal was almost constant across DS in the efficient search (F_3,1178_ = 0.12, p = 0.94), it showed a significant decrease for larger display sizes in inefficient search during the visual epoch (interaction: F_3,1190_ = 3.87, p = 0.009). The increase of the value signal in the efficient search was predominantly due to the reduction of firing to the non-preferred condition across neurons in both visual and reward epochs (Fig 3F) as can also be seen in the population average PSTHs (Fig 3B).

Critically, results showed a significant negative correlation between the magnitude of the value signal of the recorded neuron in each session and both total search time and search slope in that session in both animals (rs < -0.17, ps < 0.0001, Fig 3F). This means that larger value signal in neurons was generally concurrent with faster and more efficient value-driven visual search.

### Saccade-related firing is suppressed in TA trials in efficient search

vlPFC neural firing is not predominantly saccadic but can be similar to visuomotor responses seen in FEF (Lowe, Zinke, Cosman, & Schall, 2022). PSTHs time-locked to saccade time show perisaccadic excitation in both TP and TA trials (Supp Fig 3D-E). Nevertheless, since display and saccade onsets during the search are temporally close to each other raw PSTHs time-locked to one event contains the confounding effect of the other event. To disentangles the effect of each event on the PSTHs, a deconvolution technique can be used (Ghazizadeh, Fields, & Ambroggi, 2010). Interestingly, deconvolved responses revealed a firing ramp-up to the time of the saccade, but this ramp-up was stronger in TP vs. TA trials in the efficient search predominantly due to suppression of response in TA trials (Fig 3G). This enhancement was seen across TP- and TA-pref neurons during the saccade, suggesting that TA-pref neurons must have switched to TP preference at the time of saccade. On the other hand, there was minimal difference in deconvolved firing at the visual onset to TP and TA across the population due to the fact that TP and TA-pref neurons mostly canceled each other’s responses. While there was no significant effect of search type on the value signal in visual-related firing, this effect was significant on the value signal in saccade-related firing (F_1_,_1115_ = 18.12, p < 0.0001, Fig 3H). Once again, this saccade-related value signal decreased across the display size was significant in the inefficient search (Fig. 6C top, F_3,522_ = 5.55, p = 0.018) but not in the efficient search despite some decreasing trend (Fig. 6C bottom, F_3,593_ = 2.31, p = 0.12).

### vlPFC shows higher firing and more variable spiking during inefficient search

Search efficiency also affected the firing rate and firing variability of neurons. Overall, the firing rate in both TP and TA trials tended to be higher in inefficient compared to efficient search (Fig 4A-B, TP: F_1,2372_ = 11.74, p < 0.0001; TA: F_1,2372_ = 17.69, p < 0.0001). Firing rate was significantly higher in larger display sizes (F_3,4752_ = 5.49, p = 0.001), showing that neurons fired more when there were more objects on the screen despite the decreasing value signal (Fig 4A-B vs. Fig 3C-D). Firing rate variability can be parsed into two components one related to trial-to-trial firing rate variability (nRV) and the other related to within-trial spiking irregularity (nSI, (Fayaz, Fakharian, & Ghazizadeh, 2022)). Consistent with previous findings, nRV was reduced after display onset (Churchland et al., 2011; Fayaz et al., 2022). This transient reduction was followed by an increase later in the trial. Notably, firing rate variability in this late epoch was higher in the inefficient compared to efficient search (Fig 4C-D, TP: F_1,2330_ = 22.67, p < 0.0001). Spiking irregularity showed less fluctuation during the trial, but it too was significantly higher in inefficient search (Fig 4 E-F, TP: F_1,2330_ = 8.46, p = 0.004; TA: F_1,2328_ = 10.31, p = 0.001). Together these results suggest that the firing rate was higher and more variable in the inefficient vs efficient search.

**Fig. 4.**
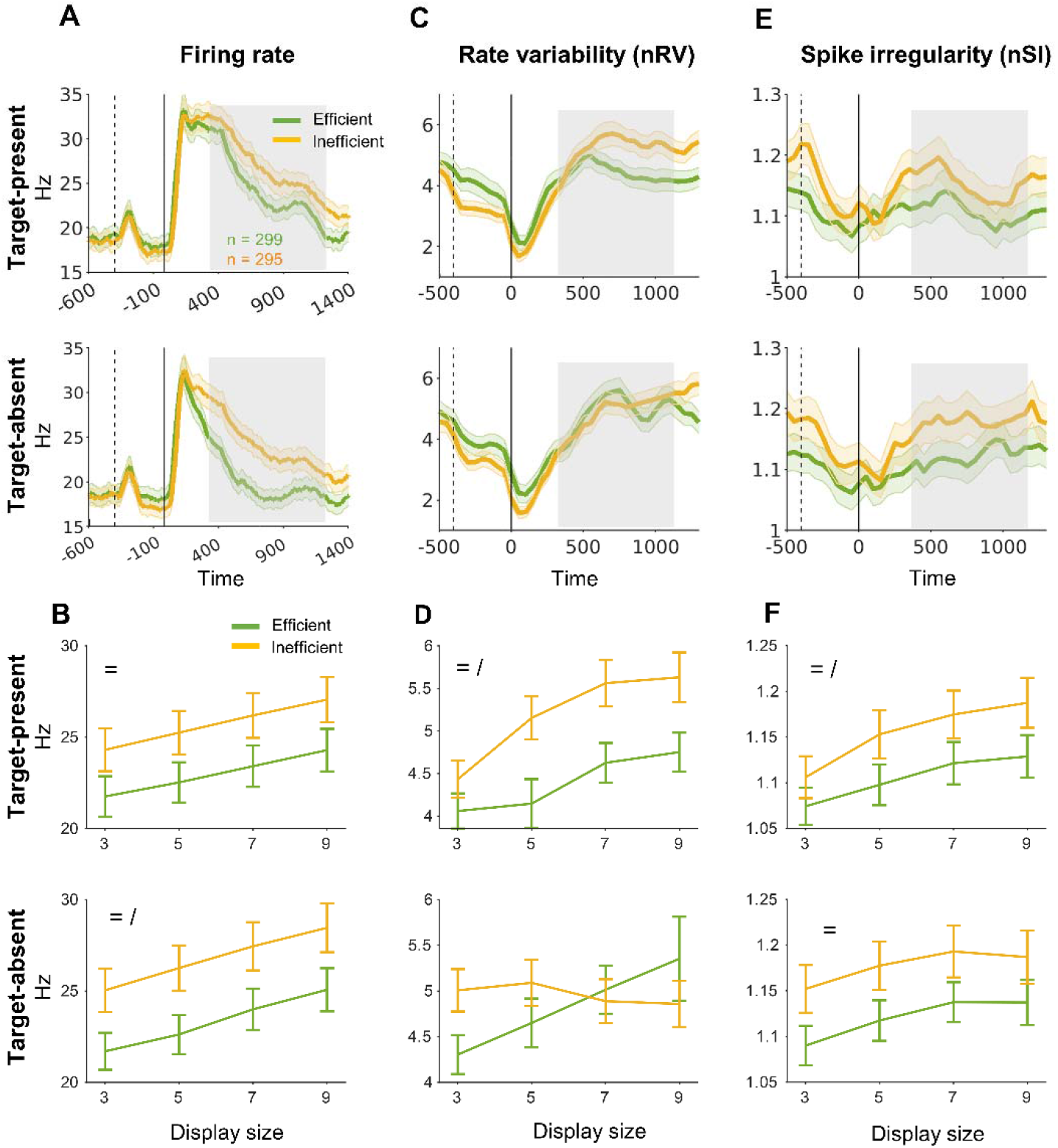
Firing rate and its variability in vlPFC neurons in inefficient and efficient searches. A) Population average of PSTH collapsed across all display sizes for the efficient (green) and inefficient (orange) searches for the TP trials (top) and the TA trials (bottom). The gray patch shows the 350-1300ms relative to the display onset used for comparing the firing rate and its variability between search types. B) The average firing for the efficient and inefficient search types in TP trials (top) and TA trials (bottom). C & D) Same format as A & B, but for between trials rate variability (nRV). E & F) Same format as A & B, but for within trial spike irregularity (nSI). (=, /, x: main effect of search efficiency, display size, and interaction, respectively; *p < 0.05, **p < 0.01, ***p < 0.001)

### vlPFC gamma-band power and duration correlate with search efficiency

Modulations in various LFP bands is thought to represent local and long-range network level involvement in the task (Brovelli et al., 2004; Buzsaki & Draguhn, 2004; Buzsáki & Wang, 2012; Kopell, Ermentrout, Whittington, & Traub, 2000). Time-frequency analysis using a continuous wavelet transform (CWT) revealed prominent modulations across LFP frequencies in the search task. There was a large increase in high gamma band (60-200 Hz) power and a concomitant decrease in beta band (12-30 Hz) power following display onset (Fig 5A). Most notably the duration of gamma was visibly longer in inefficient vs efficient search and for larger DS (Supp Fig 4). Trial by trial analysis clearly showed that the increase in power in high gamma continued to the search time (Fig 5C, top). The duration of suppression in beta band activity had a robust correlation with the reward time of each trial (Fig 5C, bottom). These results suggest the gamma band as a correlate of search time and the beta band as a correlate of upcoming reward or trial completion.

**Fig 5.**
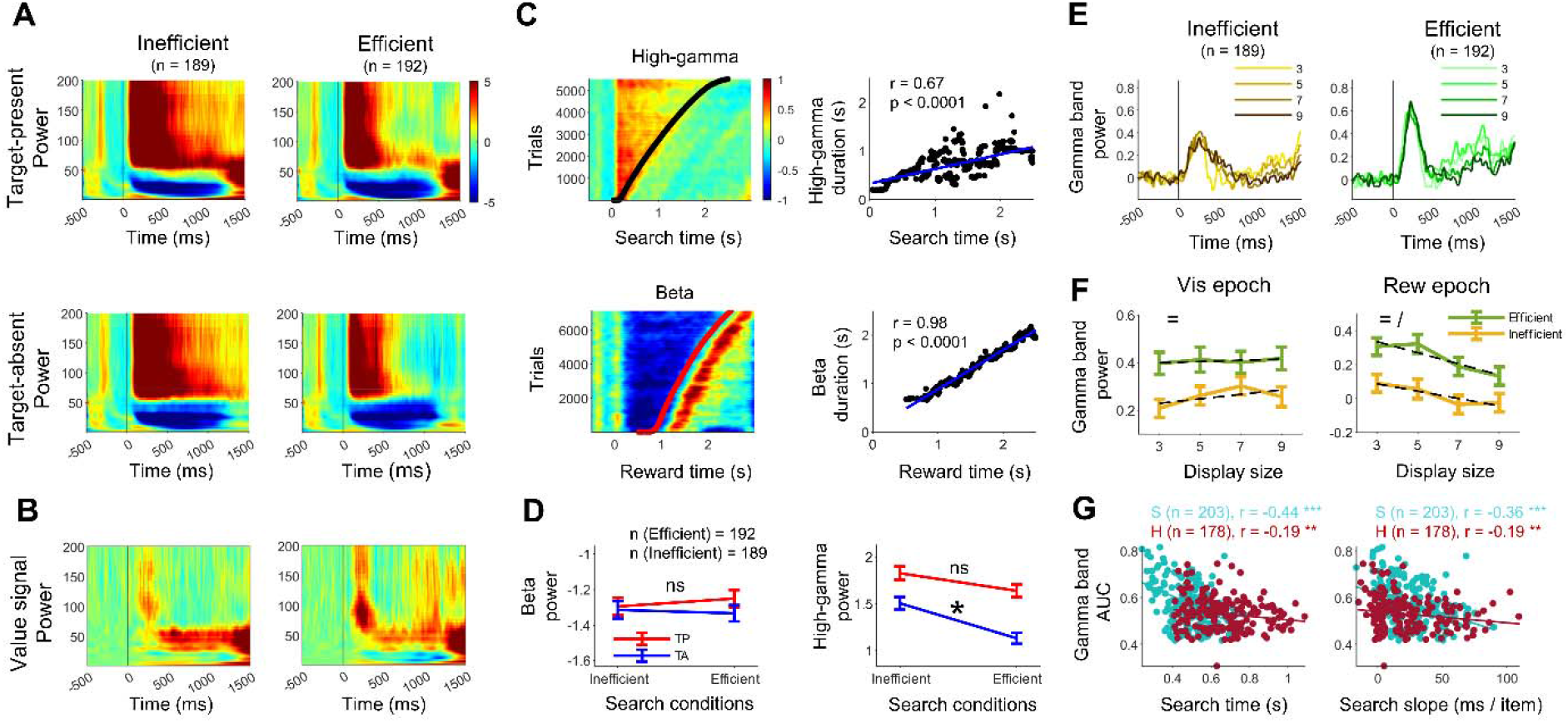
LFP and its value signal in efficient and inefficient searches. A) LFP power across time for LFP frequencies 0-200 Hz collapsed across display sizes for TP (top) and TA (bottom) in the inefficient (left) and efficient (right) searches. Colorbar shows LFP power change from average baseline power -500ms to 0 display onset. B) LPF value signal (LPF power TP minus TA) for the inefficient (left) and efficient (right) searches. C) High gamma band power sorted based on the search time across all trials. The solid black line indicates the search time (top left). Correlation of the high gamma band duration with search time across trials. Each dot is a trial. The blue line is the linear fit (top right). Beta band power sorted based on the reward delivery time across all trials. The solid red line indicates the reward time (bottom left). Correlation of the beta band duration with reward time across trials (bottom right). D) Average beta band (left) and high gamma band (right) powers for TP (red) and TA (blue) trials for the two search types. E) Value signal of gamma band power across time for all display sizes for inefficient (left) and efficient (right) searches time-locked to display onset (black solid line). F) Average gamma band value signal in visual epoch (100 to 400 ms after display onset time, left panel) and reward epoch (800 to 1100 after display onset time, right panel) across display sizes for the efficient (green) and inefficient (orange) searches. G) Scatter plots show the correlation of value AUC of high gamma band in a given session with search time (left) and search time slope (right) of TP trials separately for monkey S (cyan) and monkey H (dark red). (=, /, x: main effect of search efficiency, display size, and interaction, respectively; *p < 0.05, **p < 0.01, ***p < 0.001)

Similar to tahe firing rate, one may calculate the value signal in LFP by subtracting the powers across frequencies in TP and TA trials (TP minus TA). Results showed a robust positive value signal in the gamma range in the visual and reward epoch and a slight negative value signal in the alpha and beta range in the reward epoch (Fig 5B, Supp Fig 4-5). Notably, high gamma value signal in both visual and reward epochs was stronger in efficient vs. inefficient search (Fig. 5E-F, Fs_1,1496_ > 25, ps < 0.0001). Although the high gamma value signal was not DS dependent in the visual epoch (F_3,1496_ = 0.43, p = 0.73), it significantly decreased the reward response across display size for both searches (F_3,1496_ = 3.81, p = 0.009). For the low gamma band value signal, the effect of the search efficiency was significant only for the visual window (F_3,1496_ = 16.12, p = 0.0001), but the effect of display size was not significant for both the visual and the reward windows (ps > 0.05, Supp Fig. 5E-F). Nevertheless, the low gamma band around 500-850ms seemed to be higher in the inefficient vs. efficient search (Fig 5B, Supp Fig 4B). For the alpha and beta bands value signal, the effect of the search efficiency and the display was insignificant for the visual window (ps > 0.05), but both factors significantly affected the reward window (Fs_3,1496_ > 34.5, ps < 0.001, Supp Fig 5A-D). The positive value signal in the gamma band was due mostly to the reduction of gamma power in TA trials (Fig 5D, t_1510_ = -3.83, p = 0.0001) in the efficient search similar to what was observed in neuron firings (Fig 3F).

Finally, there was a significant negative correlation between the value signal during the visual epoch in high gamma and search efficiency across sessions (Fig 5G, rs < 0.16, ps < 0.02), similar to what was observed for neural firing value signals (Fig 3E). No significant relationship in the visual epoch was observed in alpha, beta, or low gamma bands (Supp Fig 6C & 6E). This suggests that among frequency bands, high gamma signals active search and is a good correlate of search efficiency.

## Discussion

Overtrained value can work as a strong guiding feature in visual search (Ghazizadeh et al., 2016b; Wolfe, 2020). Nevertheless, it was not previously known how the brain can use such a high-level nonphysical feature to support efficient search. Given the known role of vlPFC in visual search (Bichot et al., 2015; Bichot et al., 2019) and in value memory (Ghazizadeh et al., 2018), we recorded vlPFC neural responses in macaque monkeys while they were engaged value-driven search with various efficiencies (Fig 1). Results showed that across the population, vlPFC neurons rapidly differentiated TP from TA trials (∼150ms), and their degree of firing difference which we refer to as the value signal, were predictive of search efficiency (Fig 2-3). While in inefficient search, the value signal decreased by increasing the number of objects on the screen, display size independence emerged as the search became more efficient similar to the independence seen in behavior. The efficient search was concurrent with a lower firing rate and firing variability in vlPFC (Fig 4) and also created a differential firing when the saccade was toward the good objects (Fig 3G-H). In addition, LFP analyses revealed that among various frequency bands that showed modulations during various epochs in the trial, the gamma band oscillations significantly encoded search efficiency and marked active search time (Fig 5, Supp Fig 5-7).

The value pop-out phenomenon, which shows itself behaviorally by rapid detection of good objects by the first saccade (<300 ms), occurs for overtrained high-reward objects (Ghazizadeh et al., 2016b). Such value pop-out can be supported by activity in vlPFC, which indicated valuable target presence or absence rapidly after ∼150ms of display onset (Fig 2-3). The enhancement of value signal in the efficient search in vlPFC can play at least two roles in the task. It can indicate the presence or absence of good objects in the display as a mere perceptual signal regardless of oculomotor planning. In fact, there is evidence that vlPFC can signal the presence of good objects in a passive viewing task (Ghazizadeh et al., 2018). On the other hand, vlPFC neurons can directly promote saccade to good objects by their enhanced firing to TP vs. TA trials via their direct projections to SC (Borra, Gerbella, Rozzi, & Luppino, 2015). Interestingly, saccade-related firing in vlPFC parsed using a deconvolution technique showed enhanced differentiation in TP v TA trials in the efficient but not in inefficient search (Fig 3G).

Previous studies have shown that vlPFC region encodes and maintains the value of objects for a long time (Ghazizadeh et al., 2018). In addition to the vlPFC, certain basal ganglia nuclei, such as the caudate tail (CDt) and caudal-lateral substantia nigra reticulata and compacta (clSNr and clSNc), have robust value coding and maintain the object values for long-time (Kim, Ghazizadeh, & Hikosaka, 2015; Kim & Hikosaka, 2013; Yasuda, Yamamoto, & Hikosaka, 2012). These basal ganglia nuclei form a loop with vlPFC by sending projections to the vlPFC via the thalamus (Ghazizadeh & Hikosaka, 2021). The inferior temporal (IT) cortex also maintains the object values for a long time (Ghazizadeh et al., 2018) and has high object selectivity (Rolls & Tovee, 1995). The IT cortex is a primary input source to the vlPFC that sends the object’s visual features to the vlPFC (Gerbella, Borra, Tonelli, Rozzi, & Luppino, 2013; Webster, Bachevalier, & Ungerleider, 1994). The frontal eye field (FEF) also sends the object’s spatial information to the vlPFC (Borra et al., 2015). In addition, the vlPFC directly projects to the SC, which controls eye movements. It seems that by integrating the visual features, value memory and spatial information, vlPFC neurons are well poised to guide search behavior and send guiding information to the superior colliculus (SC) for saccade execution. In addition to the connections that have already been mentioned between the vlPFC, the basal ganglia, and the IT cortex, vlPFC also connects to other brain areas like the orbitofrontal cortex (Kennerley, Walton, Behrens, Buckley, & Rushworth, 2006), anterior cingulate cortex (Kennerley et al., 2006) and the amygdala (Paton, Belova, Morrison, & Salzman, 2006), all of which are known to mediate reward-based decisions and may play a role in value-driven search.

The LFP analyses revealed that the enhancement of gamma-band power could indicate target presence during search, with effects being significantly stronger in the efficient search (Fig. 5). While value signal in gamma band was stronger in efficient search the duration of gamma band response was longer in inefficient search similar to the higher firing of neurons in inefficient condition (Fig 4A). In particular, consistent with the previous visual search studies (Ossandón et al., 2012), the results showed that the gamma band activity was correlated with the behavioral search time (Fig 4C). This finding suggests that the gamma band activity is an index of active search (Henderson, 2003). So long as the subject searches for the target, the gamma band is active, and the neural firing is high. When the target is detected, the gamma band is deactivated. Indeed search is often thought to involve an active dimension that can be attributed to top-down frontal control (Wolfe, Butcher, Lee, & Hyle, 2003).There was also a significant beta band suppression from the display onset until the outcome delivery time, but this suppression was not predictive of search efficiency (Fig 5C-D, Supp Fig 5-6) and may better be interpreted as encoding trial termination by reward as an upcoming event consistent with some previous evidence (Roelfsema, Engel, König, & Singer, 1997).

In summary, the results of the current study revealed a candidate neural mechanism that can support value pop-out and efficient value-driven search in neural firing and LFP of vlPFC circuitry. Furthermore, our results showed a significant negative correlation between the size of the value signal in firing or gamma band power in vlPFC and the degree of search efficiency. Such quick detection of high-value objects is an object skill that can be crucial for the fitness of animals to find hidden rewards faster than their competitors (Hikosaka, Yamamoto, Yasuda, & Kim, 2013). Whether vlPFC is the first region in which this rapid value signal for differentiation of TP and TA emerges after seeing a display or is simply following the lead of other cortical and subcortical regions involved in object value memory and the causality of value signal in search efficiency remains to be tested.

## Material and Methods

### Subjects and surgery

Two rhesus macaque monkeys (*Macaca mulatta*) were used in our experiments (monkeys S and H were 12 and 10 years old, respectively). All procedures and animal care were in accordance with guidelines set by the National Institutes of Health (USA) guidelines for humane care and use of laboratory animals and were approved by the Ethical Committee of the Institute for Research in Fundamental Sciences (IPM, protocol number 99/60/1/172). A titanium head holder and a recording chamber were implanted on the head of each animal under general anesthesia during a sterile surgery in the IPM large animal operation room. The head holder was implanted on the midline of the parietal lobe for both monkeys. The recording chamber was located and laterally tilted on the right PFC for Monkey S and the left PFC for Monkey H. After the surgery, the MR images were taken from both monkeys’ heads to confirm the correct position of the recording chamber.

After the monkeys learned the experimental tasks, a second surgery was done for craniotomy over the PFC region. Grids with 1-mm spacing were placed over the chamber to record the neural data.

### Recording localization

To precisely locate the recording sites on the ventrolateral prefrontal cortex (vlPFC), T1- and T2-weighted (3T, Prisma Siemens) MR images were used. The recording chamber was filled with gadolinium before the MRI scan session to quickly locate the chamber on the monkey head in the MR images (Supp Fig. 2A). To further verify the vlPFC location in each monkey, the National Institute of Mental Health Macaque Template (NMT) toolbox (Seidlitz et al., 2018) was used to bring the D99 standard monkey brain atlas into the monkeys’ native space. This procedure enabled us to determine the accessible vlPFC region via the recording chamber for each monkey (Supp Fig. 2B).

### Stimuli

Objects with fractal geometry were used as visual stimuli (Fig. 1A, Miyashita, Higuchi, Sakai, & Masui, 1991). One fractal consisted of four point-symmetrical polygons superimposed around a common core with smaller polygons in the front. Each polygon’s properties (size, edges, color, etc.) were randomly selected. Fractal diameters averaged 4 degrees. Monkeys saw many fractals (> 600 fractals) in sets of eight either in 1 reward training session or > 5 reward training sessions (over-trained fractals) prior to their first use in visual search task.

### Task control and neural recording

All behavioral activities and recordings were controlled by custom-developed software based on C. Neural data acquisition was performed using a Cerebus Blackrock Microsystem (www.blackrockneurotech.com). Eye position was tracked with an Eyelink 1000 Plus with a sampling rate of 1000 Hz. Apple juice diluted with water (50% and 60% for monkeys S and H, respectively) was used as a reward in the experiment tasks. For each experiment session, animals were head-fixed in a monkey chair in front of a 21 inches CRT monitor (LG), which was used for visual stimuli presentation.

Tungsten epoxy-coated (FHC, 200 um thickness) or glass-coated (AlphaOmega, 250 um thickness) electrodes were used to record single-unit activity. For each recording session, the electrode was backloaded into a sharpened guide tube made of stainless steel. The dura was punctured by the guide tube, and then the electrode was inserted into the brain by the Narishige (MO-97A) oil-driven micromanipulator.

The electrode’s electric signal was digitized at 30 kHz after being amplified and filtered (1 Hz to 10 kHz). Blackrock online sorting was used to isolate the unit spike shapes. The spike time of the selected unit was sent to MATLAB to create an online peristimulus time histogram (PSTH) and raster plots for that unit. All well-isolated and visually responsive neurons were recorded and included in this study, 526 in total mostly in area 46v ventral to principal sulcus (230 Monkey S and 296 Monkey H, Supp Fig 2). These neurons were recorded during the efficient search or inefficient search or in both search types defined based on the search slope criteria presented in the result section (118 and 177 neurons in the efficient search, and 122 and 178 in inefficient search in monkeys S and H, respectively.)

### Neural data analysis

Neural spiking data were time-locked to the onset of visual (display) stimulus. The main analysis epoch was from 100 to 400 ms after visual stimulus onset. The average firing rate was computed for the Target Present (TP) and Target Absent (TA) trials within the analysis epoch. The area under the receiver operating characteristic curve (AUC) was quantified to measure the discriminability of TP and TA trials (referred to as value AUC). The Wilcoxon rank sum test was used to calculate the AUC’s statistical significance for each neuron. The preferred value for each neuron was determined using a cross-validation technique (Ghazizadeh et al., 2018).

#### Local field potential (LFP) Analysis

For the LFP analysis, data were down-sampled to 1 kHz and band-pass filtered between 0.1 to 250 Hz. Line noise and its sub-harmonics were removed using notch filters. For each session, trials with an LFP amplitude range of 3 standard deviations from the median were identified as noisy trials and removed in pre-processing. All pre-processing steps were performed using EEGLAB (-v2022.0).

The time-frequency analysis was performed using a continuous wavelet transform (CWT) and averaged over all frequencies of interest on each time sample. The CWT was obtained using an analytic Morse wavelet (Aguiar-Conraria & Soares, 2011). In this study, four frequency bands were defined as Alpha (8-12 Hz), Beta (12-30 Hz), Low-Gamma (30-60 Hz), and High-Gamma (60-200 Hz). In trial-power versus time plots (Fig 8a), CWT results were smoothed across trials and time using median and mean averaging, respectively. LFP response duration is calculated using Full Width at Half Maximum (FWHM) for each trial and correlated with the behavioral results (Fig 5).

### Onset detection procedure

To select visual responsive neurons, custom-written MATLAB functions were used as discussed before (Ghazizadeh & Hikosaka, 2021, 2022; Ghazizadeh, Hong, et al., 2018). Briefly, first, the average firing PSTH time-locked to display onset across all trials for each neuron was computed. Then, the computed firing rate was converted to z scores using the baseline from -200 to 30 ms related to the display onset. Using MATLAB findpeaks, the first response peak after object onset was detected. The minimum peak height was 1.64, corresponding to the 95% confidence interval. The first valley before the first detected peak was taken as the visual onset of the neuron. This algorithm was also used to detect the onset of the value signal by using the average value PSTH of each neuron.

### Object-value training task

To train the object values in monkeys, a biased value saccade task was used (Fig. 1B). In each trial of this task, a white dot appeared in the center of the screen for the monkey to fixate. After maintaining fixation for 200ms, a high-value (good) fractal or low-value (bad) fractal was displayed on the screen in one of the eight peripheral locations (eccentricity 9.5°). After a 400ms overlap time, central fixation disappeared, requiring the animal to saccade to the fractal and hold gaze for an additional 500 ± 100ms to receive a small (bad object) or large (good object) reward. After reward delivery, a variable inter-trial interval (ITI) of 1-1.5s was initiated with a blank screen presented. Each training session consisted of 80 trials, including 64 force trials in which each object was pseudo-randomly presented eight times and 16 choice trials in which one good and one bad fractal were displayed simultaneously and diametrically on the screen. The choice trials had the same timing structure as the force trials, but the monkey should choose one of the fractals by a saccade. If the monkey procedurally performed each trial correctly, a correct tone was played. Otherwise, an error tone was played due to an early saccade to a fractal or breaking fixation resulting in an error tone. Results in choice trials were used to measure the monkey’s knowledge of object values.

### Value-driven search task

In this task, monkeys had to find a good object surrounded by a variable number of bad objects in target-present (TP) trials or reject the trial by coming back to the center in the target-absent (TA) trials were all displayed objects were bad (Ghazizadeh et al., 2016b). Subjects learned object values before the search in the object value training task. TP and TA trials were intermixed with equal probability. Each trial began with the presentation of a fixation dot on the screen (Figure 1D). After gazing at the fixation point for 400 ms, fixation point would disappear and a display with 3, 5, 7, or 9 fractals on a 9.5°eccentricity imaginary circle would be turned on (display onset). The first (arbitrary) fractal’s angle was uniformly picked around this circle. Fractals presented in each trial were pseudo-randomly selected from a set of 24 (12 good/12 bad). The monkey would then have 3s to either choose an object by gazing at it or reject the trial by coming back to center. To choose an object, the monkey should gaze at it for 600ms (committing time). Then to get the reward it has to continue gaze for another 100ms. Breaking gaze during this 100ms would result in an error tone but the monkey was free to look away from an object before committing time has passed. Following object choice, the display would be turned off the reward corresponding to the chosen object would be delivered. A trial could be rejected either by continuing the gaze on the center dot for 900ms after display onset or by coming back to the center and gazing on the fixation dot for 300ms (Once the gaze left fixation, the fixation dot gets turned back on). The rejection of the trial led to rapid progression to the subsequent trial (after 400ms), which had a 50% chance of having a good object. The probability of receiving a reward for rejection trials was 40% for Monkey S and 60% for Monkey H. The amount of this reward was 60% of the large reward (medium reward). This was included to encourage animals to reject multiple target-absent trials in succession and to dissuade them from selecting bad fractals. After receiving the reward, there was a random ITI 1-1.5s. In each session of the search task, subjects normally performed 240 procedurally correct trials. Both monkeys had sufficient training with search task (>10 sessions) with fractals not used in this study.

### One-search and two-search neurons

During the recording of each neuron, monkeys performed the search task one or two times. The neurons that had only one search task were called single-search neurons, and the neurons with two search tasks were called two-search neurons. For almost all two-search neurons, one search task was run with over-trained fractal sets and another with sufficiently-trained fractal sets.

### Efficient and inefficient search sessions

Search sessions were divided into efficient and inefficient sessions. For this purpose, first, the search slope of the search time of TP trials (henceforth search slope) was computed across display size using the “regress” function of MATLAB for each session. Then, for one-search neurons, searches whose slope was less than percentile 40 of the search slope distribution were considered efficient sessions, and searches whose slope was higher than percentile 60 were considered inefficient sessions. For two-search neurons, if the difference of two search slopes was more 10ms per item (roughly matching with the search slope difference based on 40 and 60 percentiles), the search with a lower slope was considered efficient, and the other search was inefficient.

### Statistical tests

One-way ANOVA test was used to examine the significant effect of display size on the search time, saccade counts, accuracy, and value signal. Two-way ANOVA test was used to investigate the effect of search efficiency and display size on the search time, saccade counts, accuracy, and value signal. T-test was used to compare the neural firing rate in the efficient and the inefficient search separately for the target-present and target-absent trials. Chi-square test was used to compare the percentage of TP-preferred and TA-preferred neurons in each search type, and the percentage of nonsignificant neurons between two search types.

## Supplementary figures

**Supp Fig. 1.**
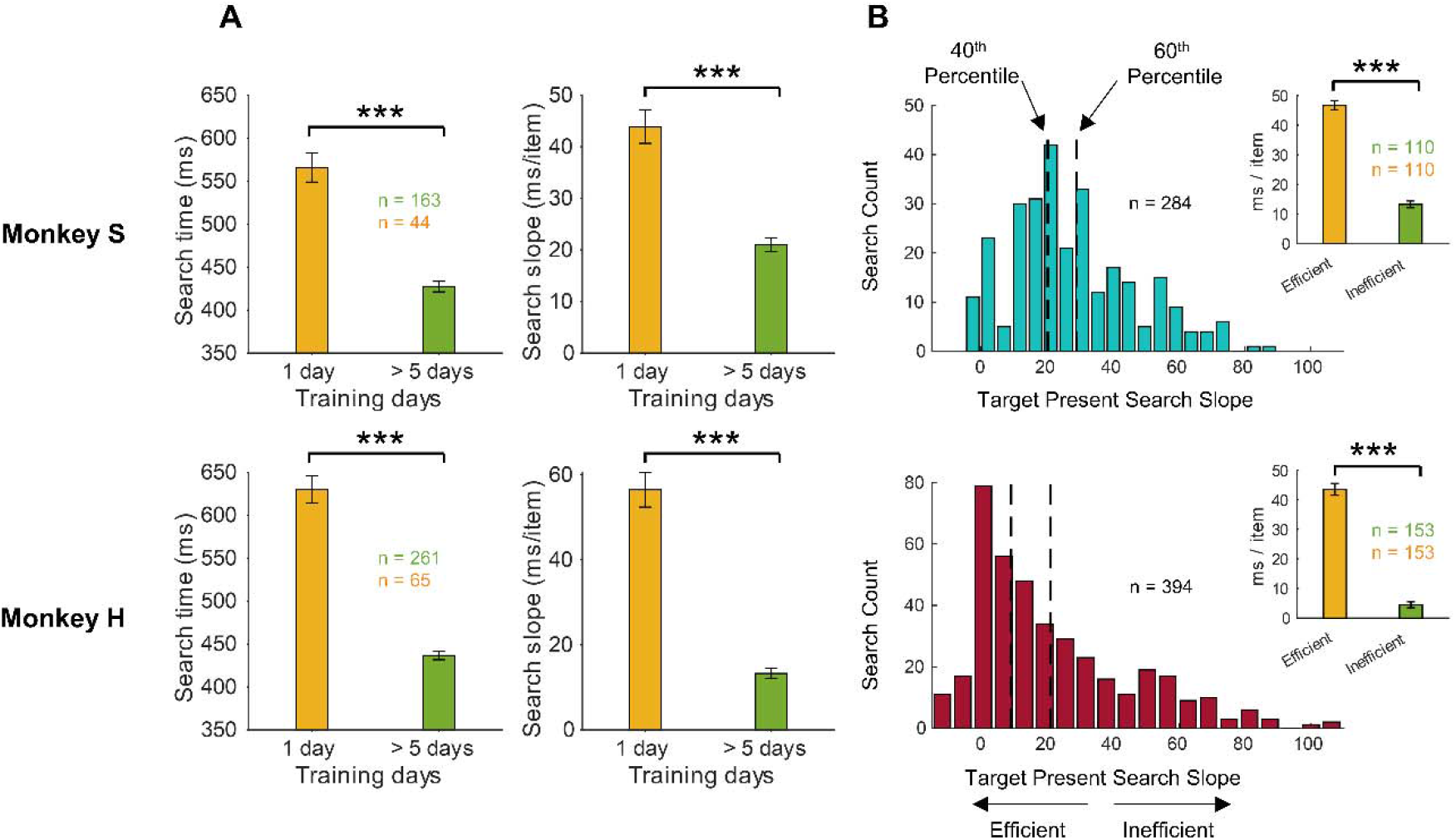
Behavior in inefficient and efficient search types. A) Search time (left) and search time slope (right) for one day trained fractal sets and >5 day trained fractal sets in TP trials for monkey S (top) and monkey H (bottom) . B) Search time slope distribution for Monkey S (top) and Monkey H (bottom) in TP trials. The x-axis shows search time slope, and the y-axis indicates the number of sessions. Sessions with a search slope lower than the 40^th^ percentile or higher than 60^th^ percentile were grouped as efficient and inefficient, respectively, for each monkey. Insets show that the average search time slope for inefficient (orange) and efficient (green) search sessions for each monkey based on the percentile definition.

**Supp Fig 2.**
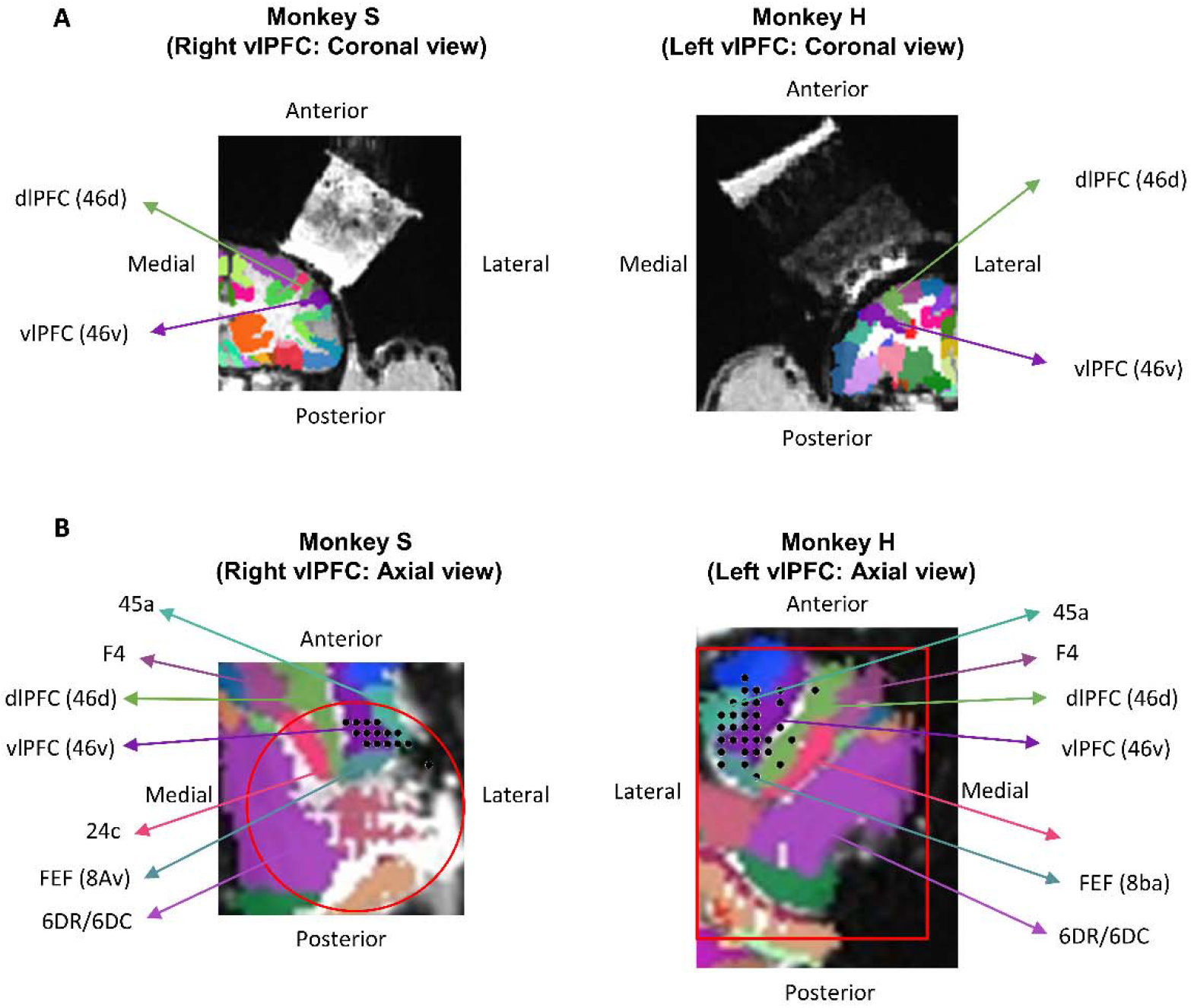
Coronal and axial views of the ventrolateral prefrontal (vlPFC) cortex and recording sites. A) Coronal view of vlPFC region (purple) of two monkeys brain registered on NMT monkey brain atlas. The recording chamber was filled with gadolinium to reveal its location on the skull for the Monkey S (left) and Monkey H (right). B) The recording sites view parallel to the chamber (right and left vlPFC for S and H monkeys, respectively) and registered on the monkey brain atlas with region numbers marked and color-coded. Each black spot shows a neural recording site. The red circle and red rectangle are the chamber locations.

**Supp Fig. 3.**
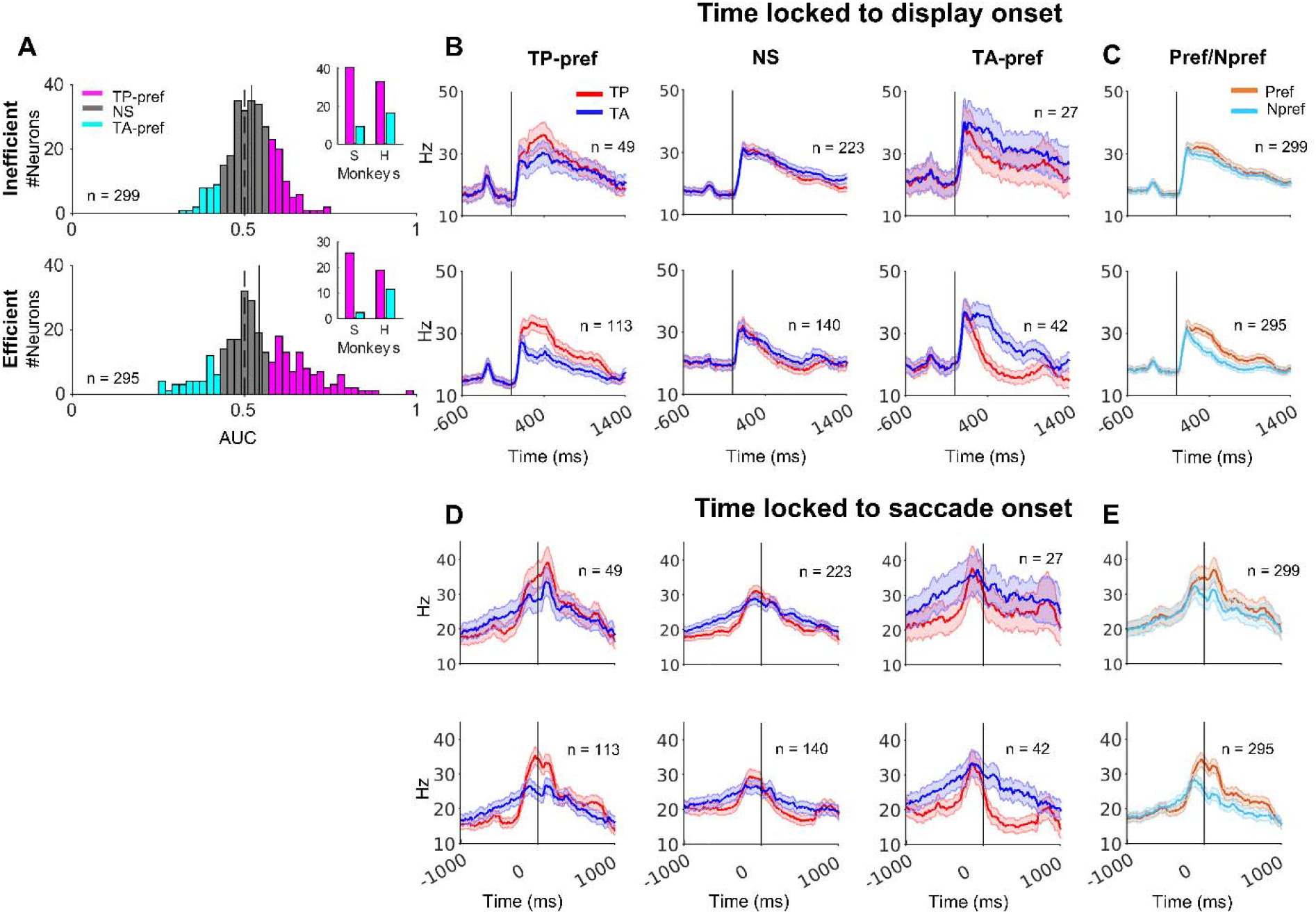
Value preference of vlPFC neurons in the inefficient and efficient searches. A) Histogram of value AUC of all recorded neurons in the efficient (top) and inefficient (bottom) search. The pink, light blue, and gray colors show the TP-preferred (TP-pref, significant AUC > 0.5), the TA-preferred (TA-pref, significant AUC < 0.5), and nonsignificant (NS) neurons, respectively. Insets indicate the percentage of TP- and TA-pref neurons for both monkeys in the search types. B) Population average of PSTH for TP-pref, NS, and TA-pref neurons in the inefficient (top) and efficient (bottom) searches. D) Population average of PSTH for preferred versus non-preferred response of neurons in inefficient (top) and efficient (bottom) searches. The neural responses are time locked to the display onset. D & E) Same format as B & C, but time locked to all saccades.

**Supp Fig. 4.**
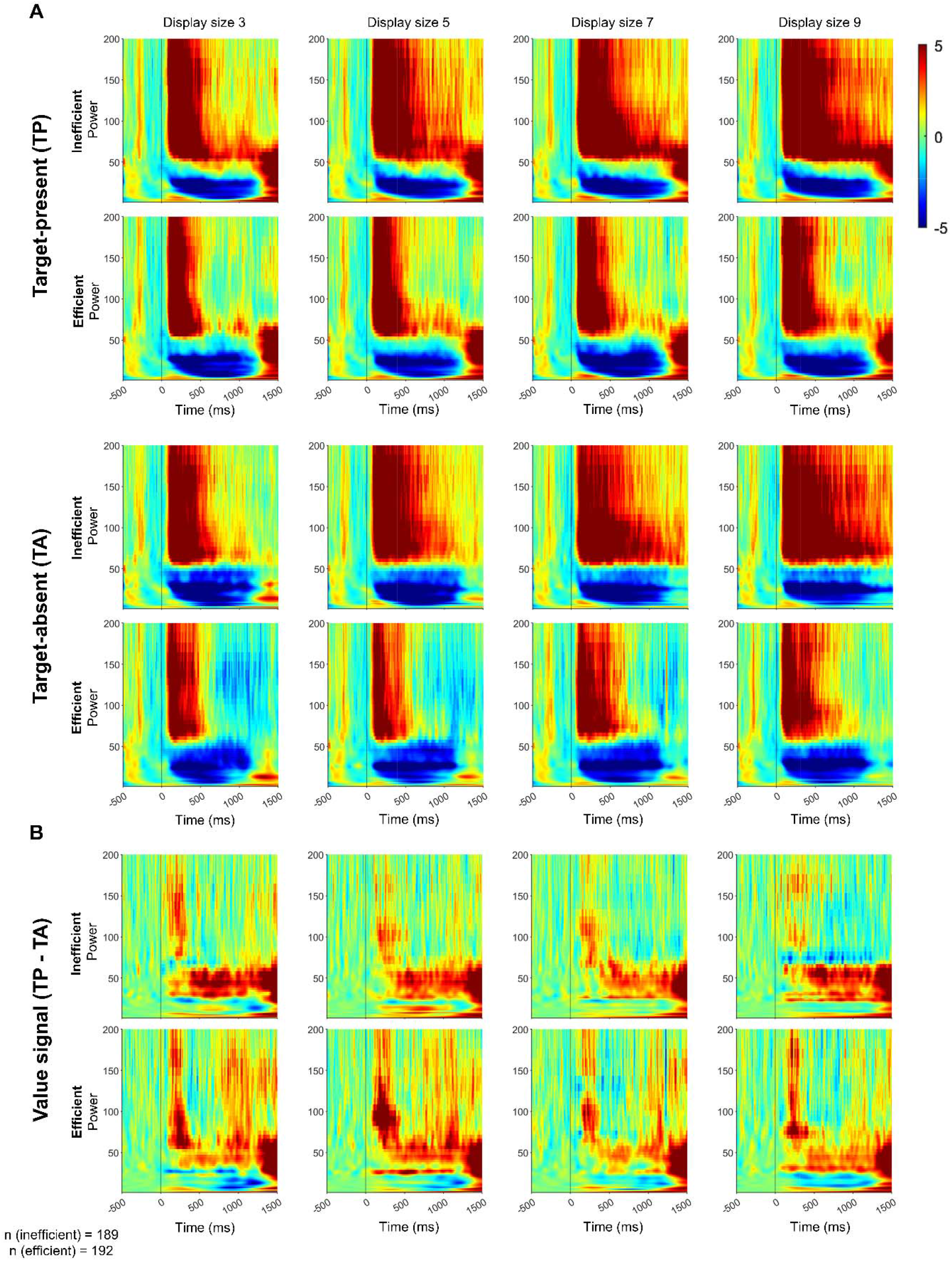
Time-frequency analysis (CWT) of LPF frequency powers across display size for inefficient and efficient searches. A) Similar format as Fig. 5A but separately for each display size. B) Similar format as Fig. 5B but separately for each display size.

**Supp Fig. 5.**
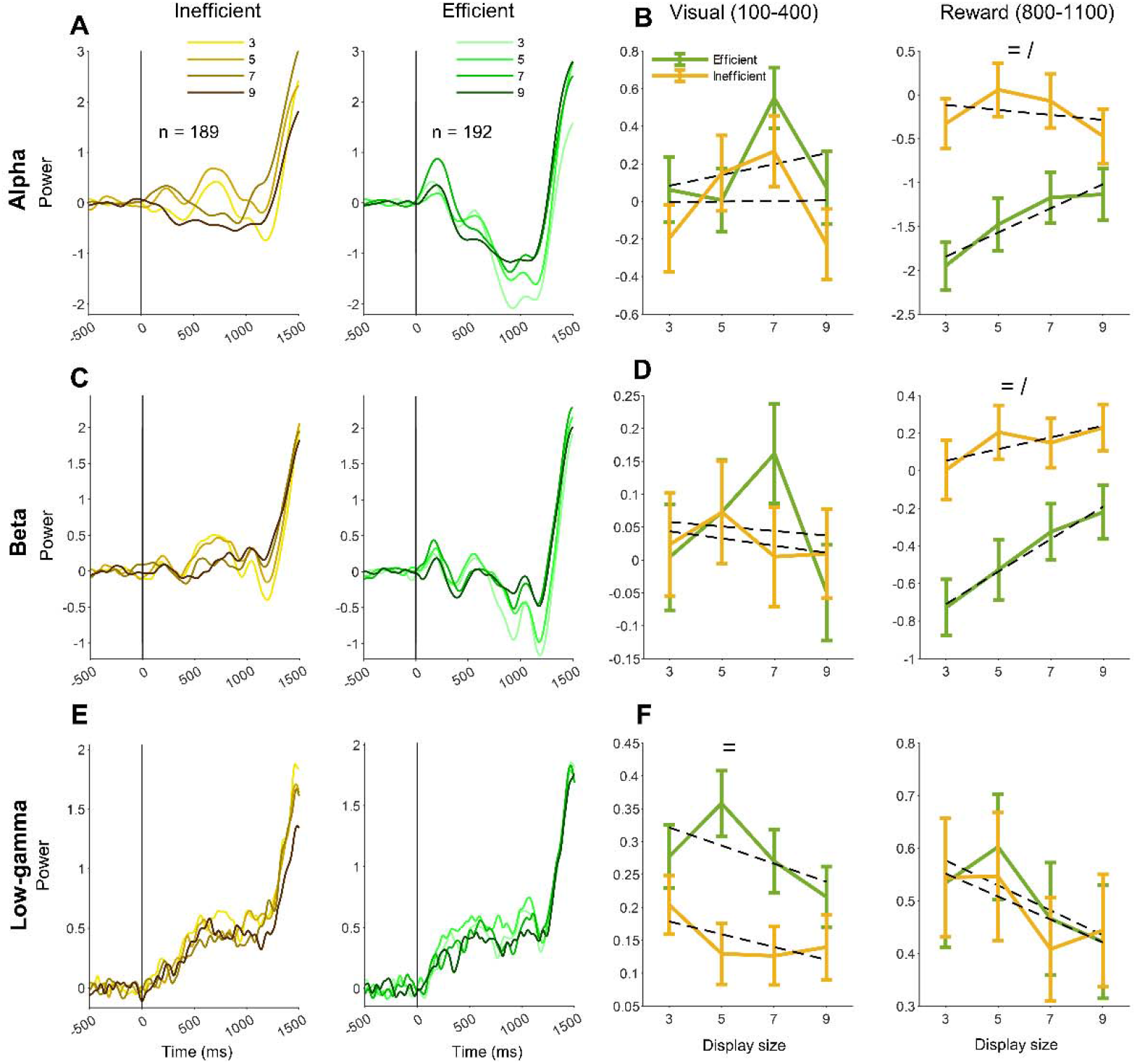
Value signal in alpha, beta and low gamma LPF frequency bands in the inefficient and efficient searches. A) Average value signal for alpha (first row), beta (second row), and low gamma (third row) bands power for different displays sizes in the inefficient (left) and efficient (right) searches LFP sessions. B) Average value signal for alpha (first row), beta (second row), and low gamma (third row) bands power across display size for inefficient search (orange) and efficient search (green) during visual epoch (100-400 relative to display onset time, left panel) and reward epoch (800-1100 relative to display onset time, right panel). (=, /, x: main effect of search efficiency, display size, and interaction, respectively; *p < 0.05, **p < 0.01, ***p < 0.001)

**Supp Fig. 6.**
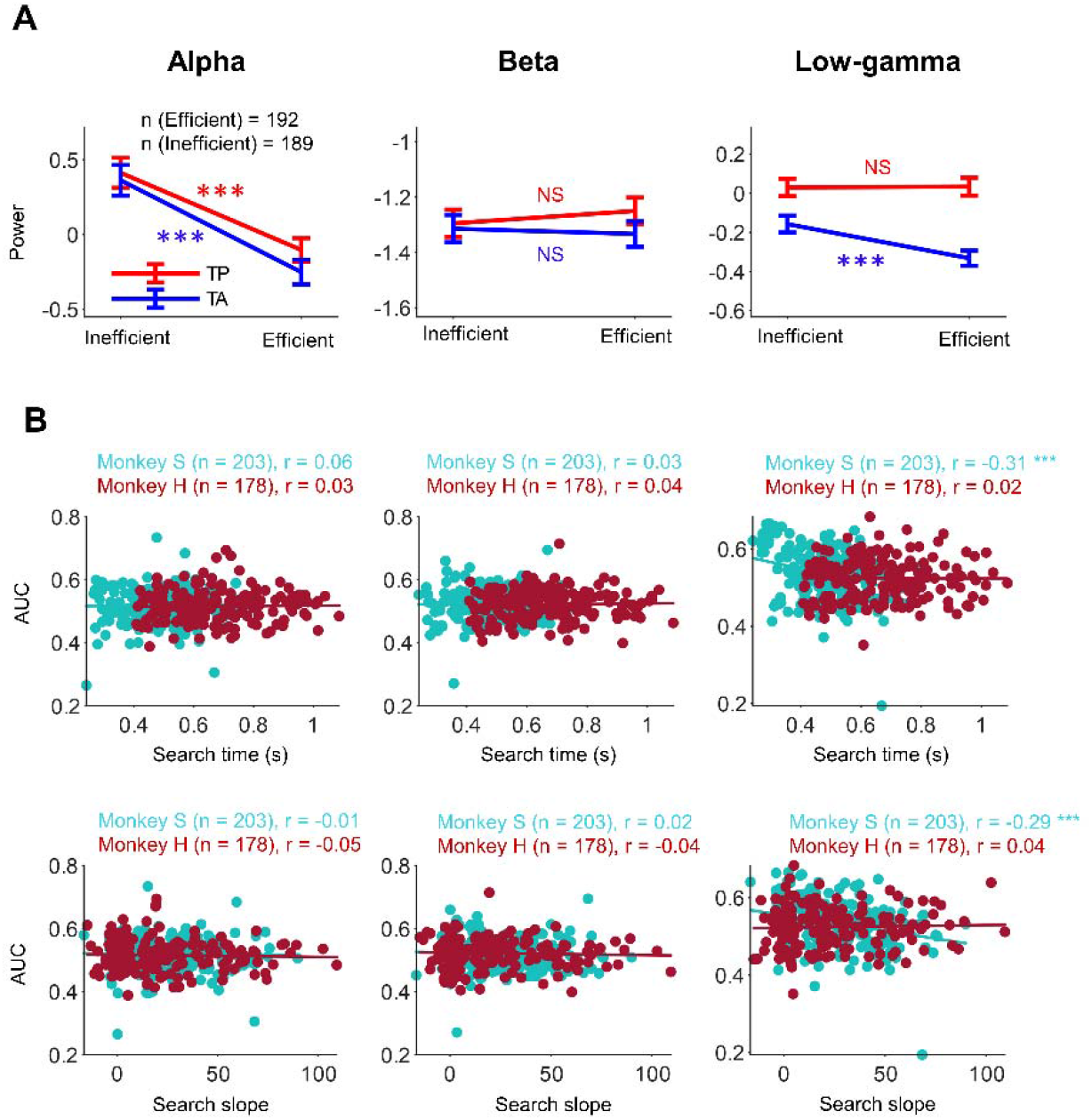
Search efficiency and alpha, beta and low gamma LFP frequency bands. A) Same format as Fig 5D but for alpha, beta and low gamma LFP frequency bands. B) Similar format as Fig 5G but for alpha, beta and low gamma LFP frequency bands. (*p < 0.05, **p < 0.01, ***p < 0.001)

## Notes

### Competing Interest Statement

The authors have declared no competing interest.

### Summary of Updates

We noticed in the current version of the manuscript that second author's first name need to be corrected. Correct name is "Armin", but we wrote "Amin". I corrected this in the new version of manuscript.

